# Using Ancient Samples in Projection Analysis

**DOI:** 10.1101/025015

**Authors:** Melinda A. Yang, Montgomery Slatkin

**Affiliations:** Department of Integrative Biology, University of California, Berkeley, CA 94720-3140

**Keywords:** projection, human population genetics, human demography, ancient genomes

## Abstract

Projection analysis is a useful tool for understanding the relationship of two populations. It compares a test genome to a set of genomes from a reference population. The projection’s shape depends on the historical relationship of the test genome’s population to the reference population. Here, we explore the effects on the projection when ancient samples are included in the analysis. First, we conduct a series of simulations in which the ancient sample is directly ancestral to a present-day population (one-population model) or the ancient sample is ancestral to a sister population that diverged before the time of sampling (two-population model). We find that there are characteristic differences between the projections for the one-population and two-population models, which indicate that the projection can be used to determine whether a test genome is directly ancestral to a present day population or not. Second, we compute projections for several published ancient genomes. We compare three Neanderthals, the Denisovan and three ancient human genomes to European, Han Chinese and Yoruba reference panels. We use a previously constructed demographic model and insert these seven ancient genomes and assess how well the observed projections are recovered.

## INTRODUCTION

The projection of a test genome onto a reference panel provides insight about the demographic relationship between the test population from which the test genome is sampled and the reference population (Yang et al. 2014). The projection shows the probability of observing a derived allele at a particular site in a test genome, relative to the derived allele frequency of the reference population for that site. Thus, using a test genome that is a member of the reference population would give a projection of one for all derived allele frequency categories. If the test genome does not belong to the reference population, then the projection may show that the test genome has more or fewer derived alleles than expected given the derived allele frequency in the reference panel.

Yang et al. (2014) showed that for a two-population scenario with no migration or population size changes, if the reference panel was sampled from one population and a test genome from the other, the projection is dependant on the effective population size and the time of divergence between the two populations. The projection is given by 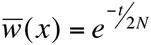, where 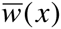 is the projection, *x* is the derived allele frequency in the reference panel, *t* is the time of divergence and *N* is the effective population size. As the two populations diverge further back in time, it is less likely to find a derived allele found in the reference panel in the test genome. A small amount of past migration from the reference population into the test population has little effect on the projection. Migration from the test into the reference population, however, increases the projection for small *x*, indicating more low frequency derived alleles are found in the test genome than expected. Population size changes, particularly in the reference population, also alter the projection such that the number of derived alleles in the test genome for different derived allele frequency categories varies with *x*. The two demographic processes that have the greatest effect on the shape of the projection are population size changes in the reference population and admixture from the test population into the reference population (Yang et al. 2014).

Here, we explore how the projection of an ancient sample depends on the relationship to present-day populations. Then, we present the projections of several ancient hominin genomes onto present-day human populations, as represented by Phase 3 of the 1000 Genomes (1KG) Panel (The 1000 Genomes Project Consortium 2012).

## SIMULATIONS OF ANCIENT SAMPLES

To simulate demographic scenarios including ancient samples, we used *fastsimcoal2* (version 2.1, Excoffier et al. 2013) to model several demographic histories, from which samples were taken to form a reference panel of *n* = 200 and a test genome to project onto the reference panel. For each simulation, we projected an ancient sample onto a modern population or a modern sample onto an ancient population. The ancient samples were taken at 500, 1000, 2000, 3000 and 4000 generations ago (ga). Unless otherwise indicated, the effective population size was 5000. We considered two demographic models: a one-population model (OPM, Figure 1, OPM A-E) where the ancient sample was directly ancestral to the present day population, and a two-population model (TPM, Figure 1, TPM A-E) where the ancient sample belongs to a sister population that diverged from the present day population.

**Figure 1:**
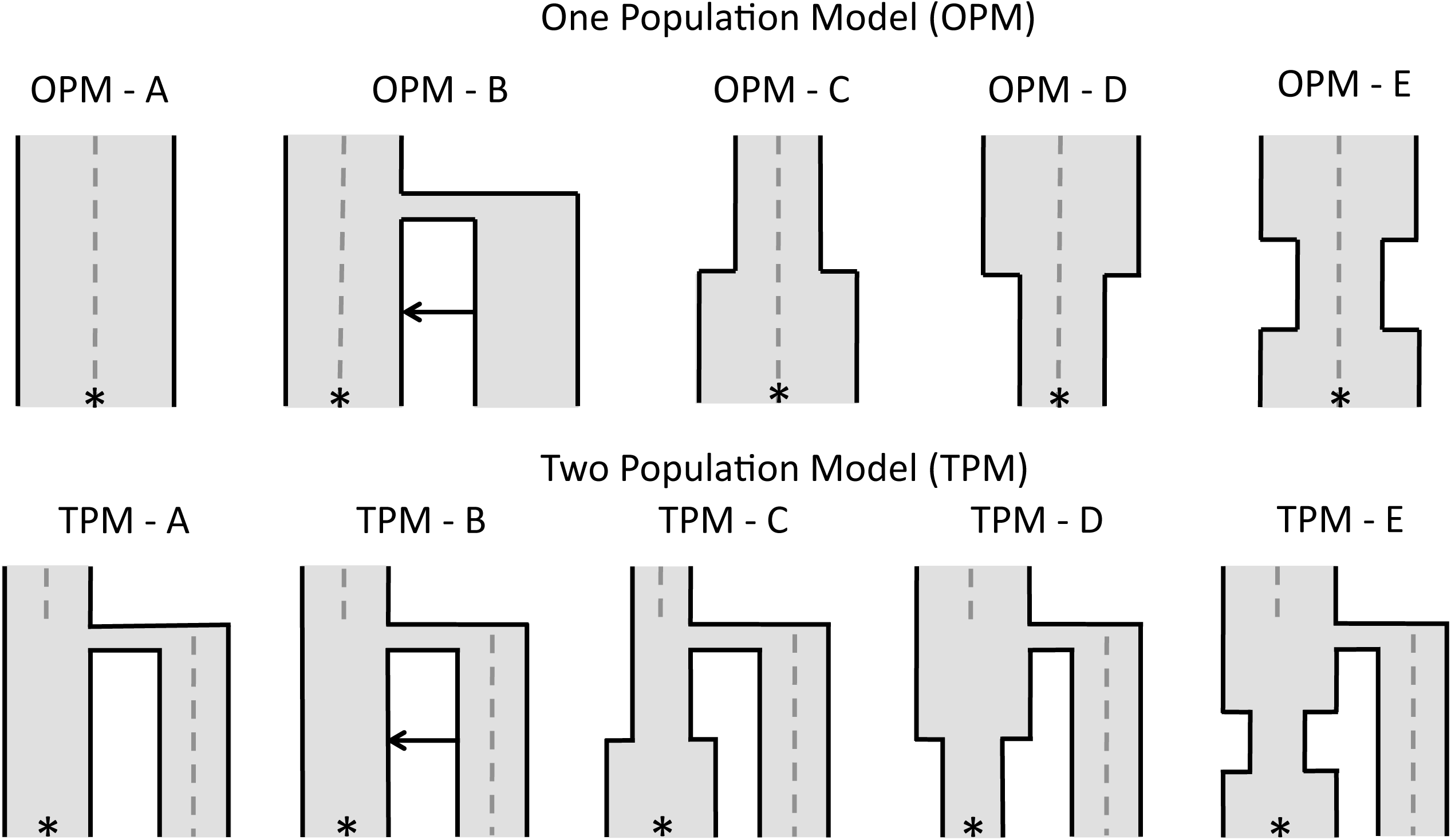
Simulated demographic models used to illustrate the effect of ancient samples in a one-population and two-population model. The * represents where the present day population was sampled and the gray dashed line indicates when the ancient genomes were sampled (0 - 4k gen). Any divergence occurs 2k gen ago. For both OPM and TPM, A has an N_e_of 5k, with no population size changes or admixture. B adds a pulse of admixture from the second diverging population. C has no admixture but allows a population size expansion from 500 to 5k in the reference population 750 gen ago. D allows the reverse, a population size decline from 5k to 500 in the reference population 750 gen ago. E has a bottleneck from 5k to 500, 500-1000 gen ago. Any diverging population has the same N_e_ as the ancestral population.

In OPM A, no population size change or migration was applied to the population. In OPM B, we applied a pulse of admixture of 0.05 at 750 ga from an unsampled population into the present-day population. We then allowed a population size expansion from 500 to 5,000 at 750 ga (OPM C), a population size decline from 5,000 to 500 at 750 ga (OPM D), and a bottleneck 500 to 1,000 ga, where the population reduces from 5,000 to 500, before recovering to 5,000 (Figure 1, OPM C-E). In the two-population model, the five same scenarios were simulated. Again, we considered no population size changes or migration (TPM A), before adding migration from the sister population into the present day population (TPM B). The three population size changes occur only in the present-day population (Figure 1, TPM C-E).

For the one-population model, when the reference panel is from the present and the test genome is ancient, the projection’s shape does depend on the sampling time (Figure 2, top row). In Figure 2, the projection of an ancient sample onto a reference panel comprised of members of the descendant population decreases with the age of the sample. With no population size changes or migration (Figure 2, OPM A), the projection follows the 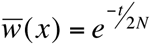 line, where here, *t* is the age of the ancient sample, not the time of population divergence. Small amounts of admixture from an unsampled population have no effect on the projection (Figure 2, OPM B). Population size changes show different levels of effect for different sampling times. When there is a population expansion, the projection decreases for rare alleles (Figure 2, OPM C), while when there is a population decline, the projection increases for rare alleles (Figure 2, OPM D). A bottleneck results in a humped shape similar to that observed when the test genome is sampled from a related population that diverged prior to the bottleneck (Figure 2, OPM E). Changes in the sampling time results in slight changes in the shape of the projection, but retains the characteristic shape for that type of population size change.

**Figure 2:**
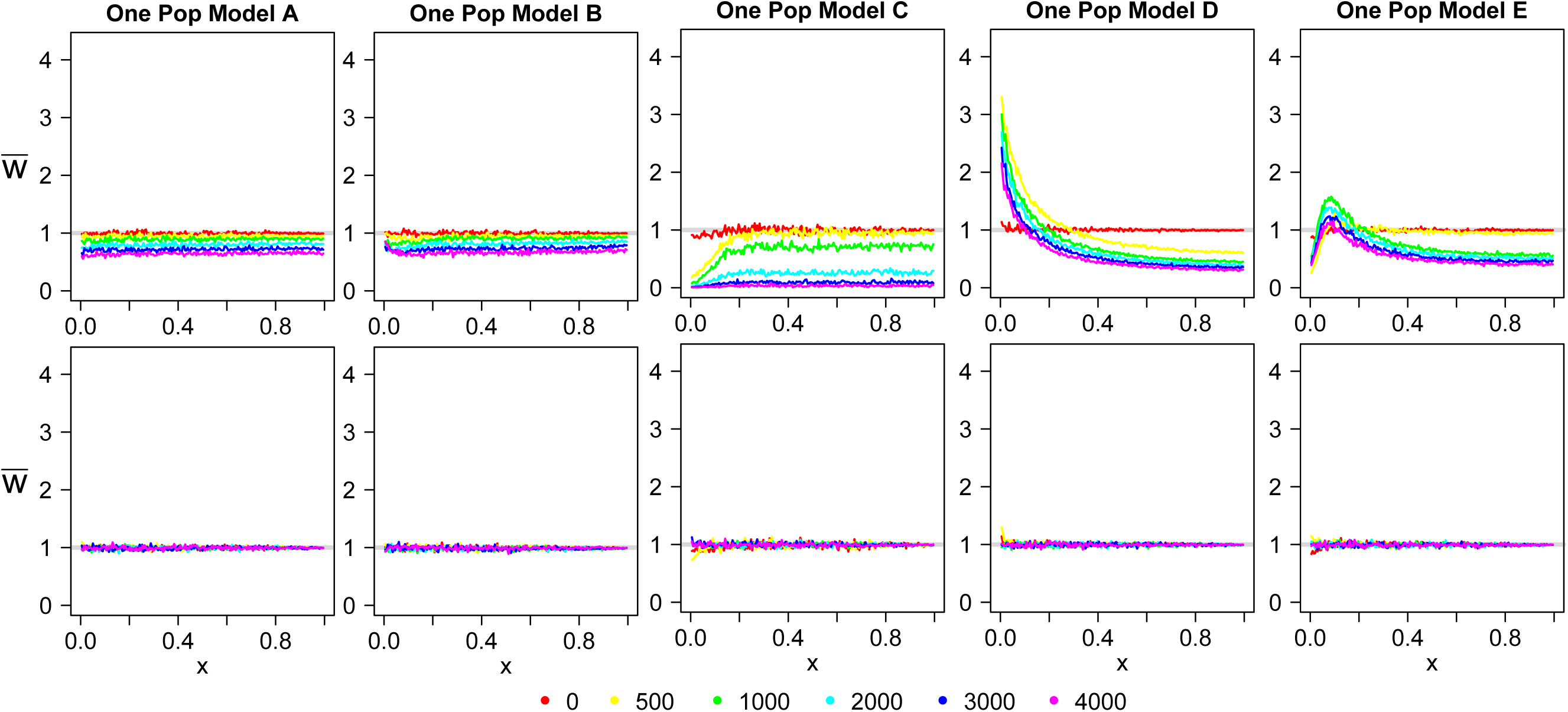
One Population Model (OPM) simulated projections for the demographic models tested in Figure 1. The key indicates the time the ancient genomes were sampled. The top row gives the results for when the reference panel is sampled from the present and the test genome is sampled from the past. The bottom row gives the reverse. 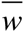 is the value of the projection and *x* is the derived allele frequency in the reference population.

The mirror scenario, where the reference panel consists of ancient samples and the test genome is sampled from the present, looks markedly different (Figure 2, bottom row). Here, the present-day test genome looks no different from the ancient population upon which it is projected. This is reasonable because the main contribution to deviations in the projection from 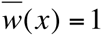is new mutations in the reference population that are not found in the test population. When the reference panel is made up of ancient samples, there are no new mutations in the reference population that are not also present in the present-day population from which the test genome is sampled. Thus, using an ancient reference panel and a test genome from the descendant population will not give insight into the demographic changes that the population has undergone between the time of sampling and the present day.

In the two-population model, the results for the projection are very different than that found for the one-population model. The simplest scenario (Figure 3, TPM A) highlights a difference in the projection relative to OPM A (Figure 2). In TPM A, the projection is lower for ancient samples, until the time of sampling is younger than the time of divergence. When the time of sampling is younger than the time of divergence, the projection no longer changes as the sampling time changes — it looks the same as if the test genome was sampled from the present day. Thus, if the time of sampling is known, the projection can determine whether an ancient sample is directly ancestral to a present day population or a member of a related population that diverged before the time of sampling.

**Figure 3:**
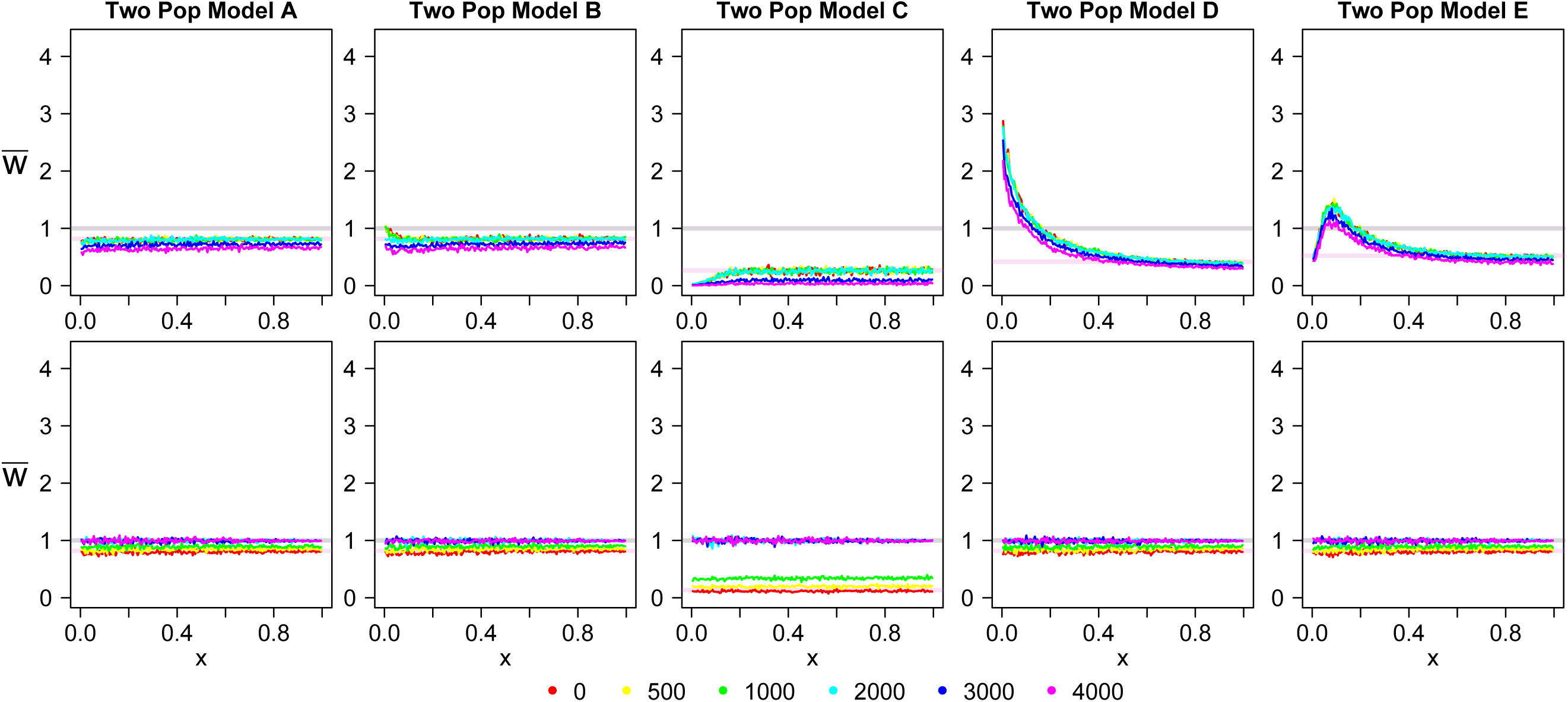
Two Population Model (TPM) simulated projections for the demographic models tested in Figure 1. The key indicates the time the ancient genomes were sampled. The top row gives the results for when the reference panel is sampled from the present and the test genome is sampled from the past. The bottom row gives the reverse. *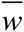* is the value of the projection and *x* is the derived allele frequency in the reference population.

A pulse of admixture from the test population into the reference population shows an increase in rare alleles, but only if the test genome was sampled after the time of divergence (Figure 3, TPM B). Population size changes show the characteristic effects (decline in rare alleles for population expansion; increase in rare alleles for population decline; ‘humped’ effect for population bottleneck; Figure 3, TPM C-E). Similar to the TPM A case, the projections for test genomes sampled more recently than the time of divergence look the same as for when the test genome was sampled in the present.

In the two-population model, when the reference panel consists of ancient samples and the test genome is sampled from the present day, the projection is again different than the reverse (Figure 3, bottom row). As the reference panel is sampled closer to the time of divergence, the projection moves closer to the 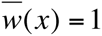 line and away from the 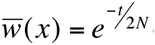 expected if the reference panel was sampled from the present. Once the reference panel is sampled from a time at least as old as the time of divergence, the projection acts similarly as in OPM A; the test genome looks as if it was sampled from the reference population — that is, 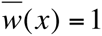for all *x*.

To conclude, the shape of the projection can be affected by the time of sampling. Particularly, the dynamics are notably different when the ancient samples are directly ancestral to the present day samples and when they belong to a sister population that diverged from the present-day population. In the following analysis, we highlight when this distinction can be made with ancient hominin data.

## PROJECTIONS OF NEANDERTHALS, DENISOVANS, AND OTHER HUMANS

Seven ancient genomes were compared to present-day human populations using projection analysis. Of the seven, three are Neanderthal, one is the Denisovan genome and three are ancient modern humans. Table 1 indicates the sampling time, as indicated by the study in which the genome was sequenced. The Altai Neanderthal (Prüfer et al. 2014) and the Denisovan (Meyer et al. 2012, Reich et al. 2010) were presented in Yang et al. (2014), and are included here for comparison with the other ancient genomes. The Vindija Neanderthal was the original Neanderthal genome sequenced (Green et al. 2010), and the Mezmaiskaya Neanderthal was sequenced by PrÜfer et al. (2014).

**Table 1:** Data used for each ancient genome.

| **Sample** | **Date Used<sup>a</sup>** | **Covg<sup>b</sup>** | **Reference** | **MinCov<sup>c</sup>** | **MaxCov<sup>c</sup>** |
| --- | --- | --- | --- | --- | --- |
| Denisovan | 75000 | 30 | Meyer et al. 2012 | 15 | 47 |
| Altai | 50000 | 52 | Prüfer et al. 2014 | 30 | 81 |
| Vindija | 40000 | 1.3 | Green et al. 2010 | 1 | 4 |
| Mezmaiskaya | 65000 | 0.5 | Prüfer et al. 2014 | 1 | 3 |
| Ust-Ishim | 45000 | 42 | Fu et al. 2014 | 21 | 64 |
| Loschbour | 6000 | 22 | Lazaridis et al. 2014 | 5 | 25 |
| Stuttgart | 5000 | 19 | Lazaridis et al. 2014 | 7 | 34 |

<sup>a</sup> in thousands of years, roughly taken from the date ranges found in the reference.
<sup>b</sup> the average coverage given in the reference.
<sup>c</sup> the 2.5% and 97.5% interval cutoffs for the coverage that were used in the analysis.

The three ancient modern humans used in this study are the Ust-Ishim (Fu et al. 2013), the Loschbour and the Stuttgart genomes (Lazaridis et al. 2014). The Ust-Ishim was sampled 45 kya, and is found to be equally distant from all present-day non-Africans, with some greater admixture into present day East Asians (Fu et al. 2013). The Loschbour and Stuttgart genomes date to around 7-8 kya, in Central Europe. The Loschbour genome was found near hunter-gather sites, while the Stuttgart genome was found with the Linearbandkeramik farming culture. Both of these genomes are of West Eurasian ancestry and are members of populations that contributed to present day European populations (Lazaridis et al. 2014).

We project these seven genomes onto three reference panels representing Europeans (CEU), Han Chinese (CHB) and the Yoruba (YRI) populations. To calculate the projection, we modified the analysis from that found in Yang et al. (2014) to use reads instead of genotypes called from the reads, in order to more accurately assess low coverage samples. We used the CEU, CHB and YRI panels from Phase 3 of the 1000 Genomes Project as the reference panels (The 1000 Genomes Project Consortium 2012). We considered only biallelic sites where the mutation was a transversion. We filtered out any sites where the mapping quality was less than 30, and for each ancient genome we filtered for sites where the coverage was within the 2.5% to 97.5% interval of the coverage distribution unique to each sample (Table 1, minCov and maxCov). The derived allele frequency of the reference panel was determined by using the genotypes assessed in the Phase 3 panels and the ancestral allele called in the Phase 3 1000 Genomes data set. For each site, the test genome was called derived or ancestral by choosing randomly from the set of reads for that site. The projection was calculated across all autosomal sites that were not filtered out by the above criteria. A minimum projection value (MPV) was calculated using the average projection for *x* > 0.5.

In the projections, there are several notable characteristics (Figure 4-6, black curve). First, with respect to the reference panel refCEU (Figure 4), the projections for the ancient samples can be divided into three main groups. The Neanderthals and Denisovan have the lowest projections, with a mean MPV of 0.4579 (sd = 0.0171). All three Neanderthals show a substantial increase in rare alleles, while the Denisovan projection shows an increase in rare alleles, but less pronounced than in Neanderthals. The Ust-Ishim shows the next lowest MPV of 0.9027 with minor deviances from a horizontal line likely indicative of population size changes in the refCEU population. The Loschbour and Stuttgart genomes sit almost entirely on the *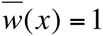*line, with a slight decrease in rare alleles.

**Figure 4:**
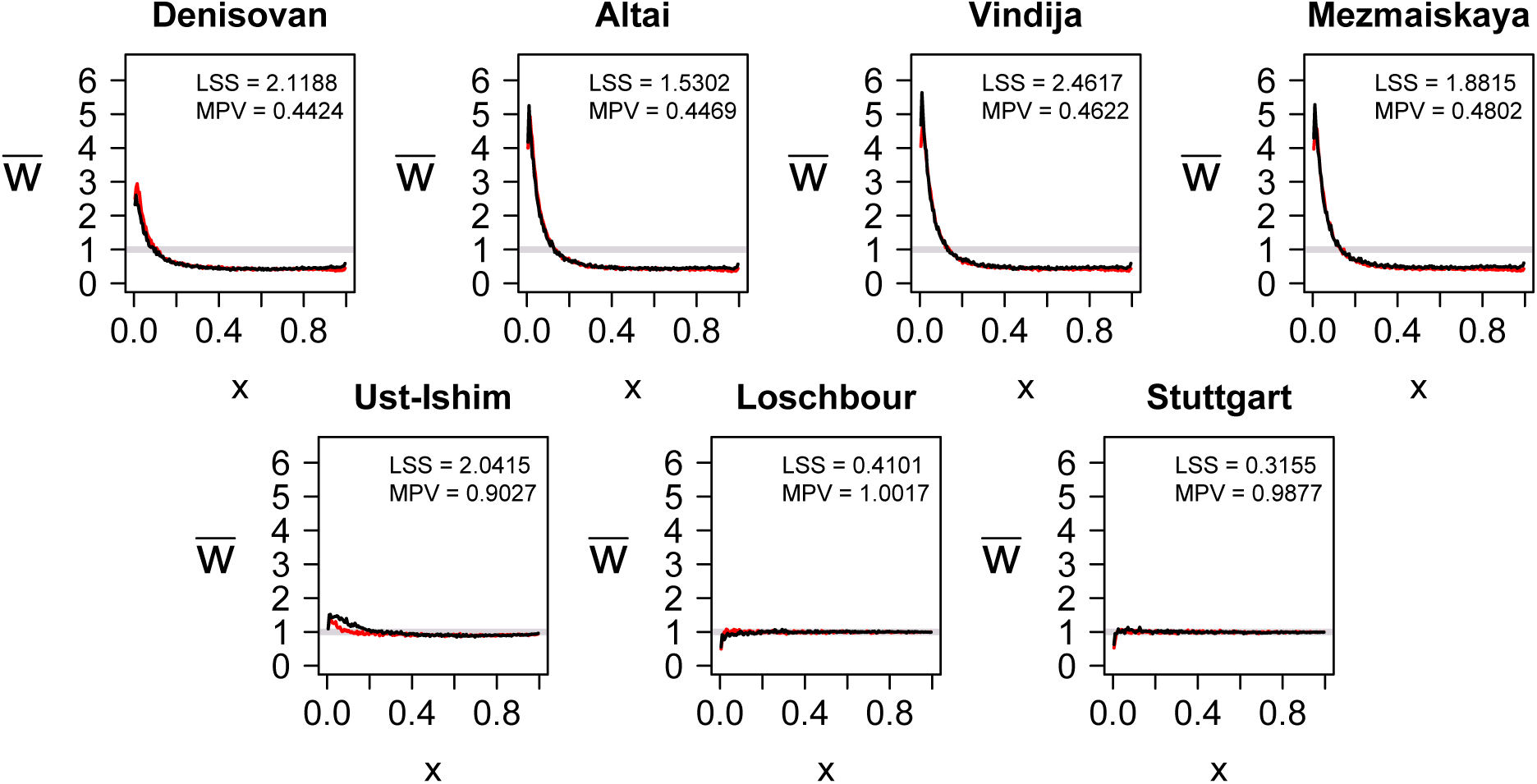
Projection of ancient hominin genomes onto the European reference panel, refCEU (black), and simulated projection (red). The sum of least squares (LSS) score gives the fit between the observed and simulated projections. The mean projection value (MPV) is the mean for *x* > 0.5.

**Figure 5:**
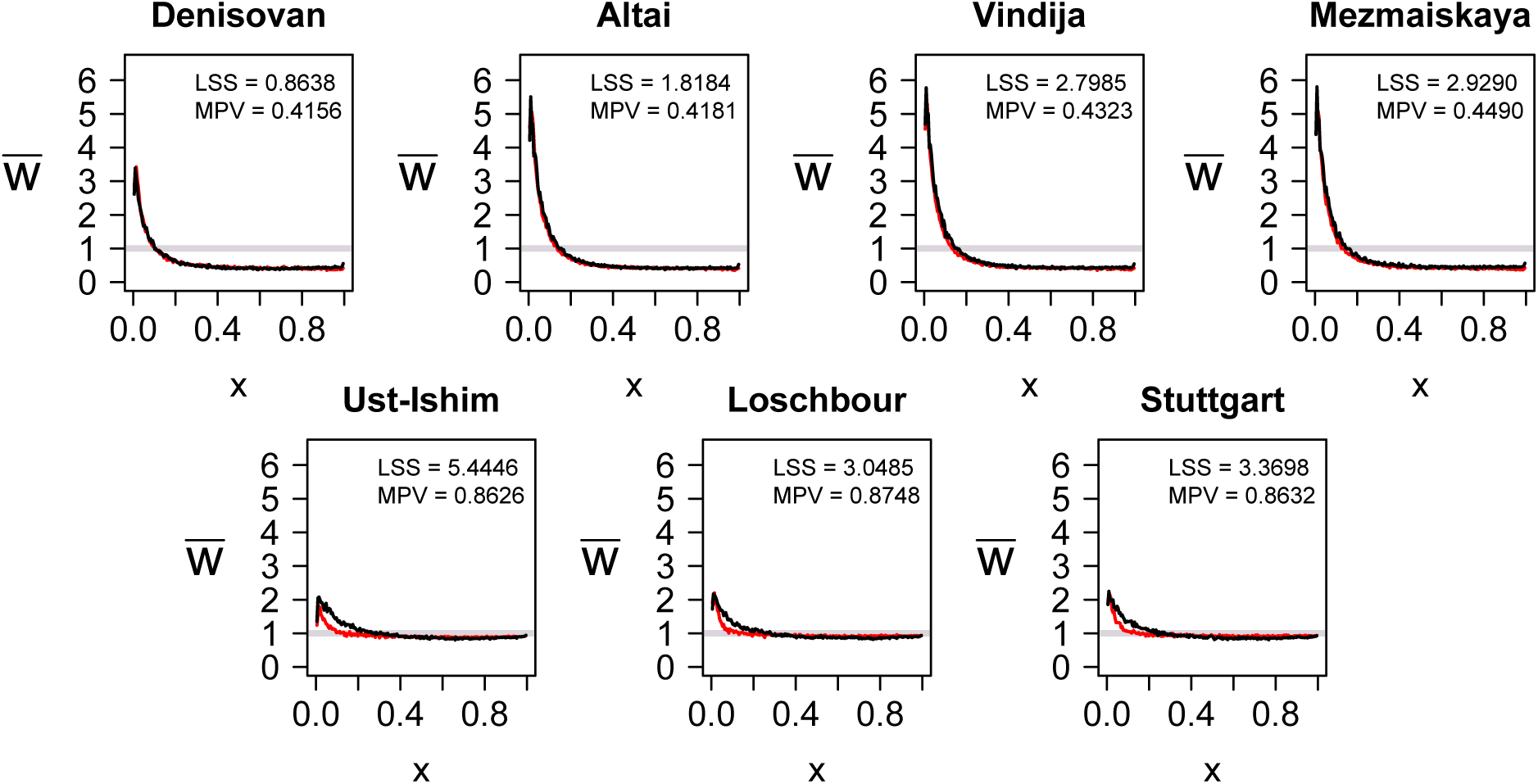
Projection of ancient hominin genomes onto the Han Chinese reference panel, refCHB (black), and simulated projection (red). The sum of least squares (LSS) score gives the fit between the observed and simulated projections. The mean projection value (MPV) is the mean for *x* > 0.5.

**Figure 6:**
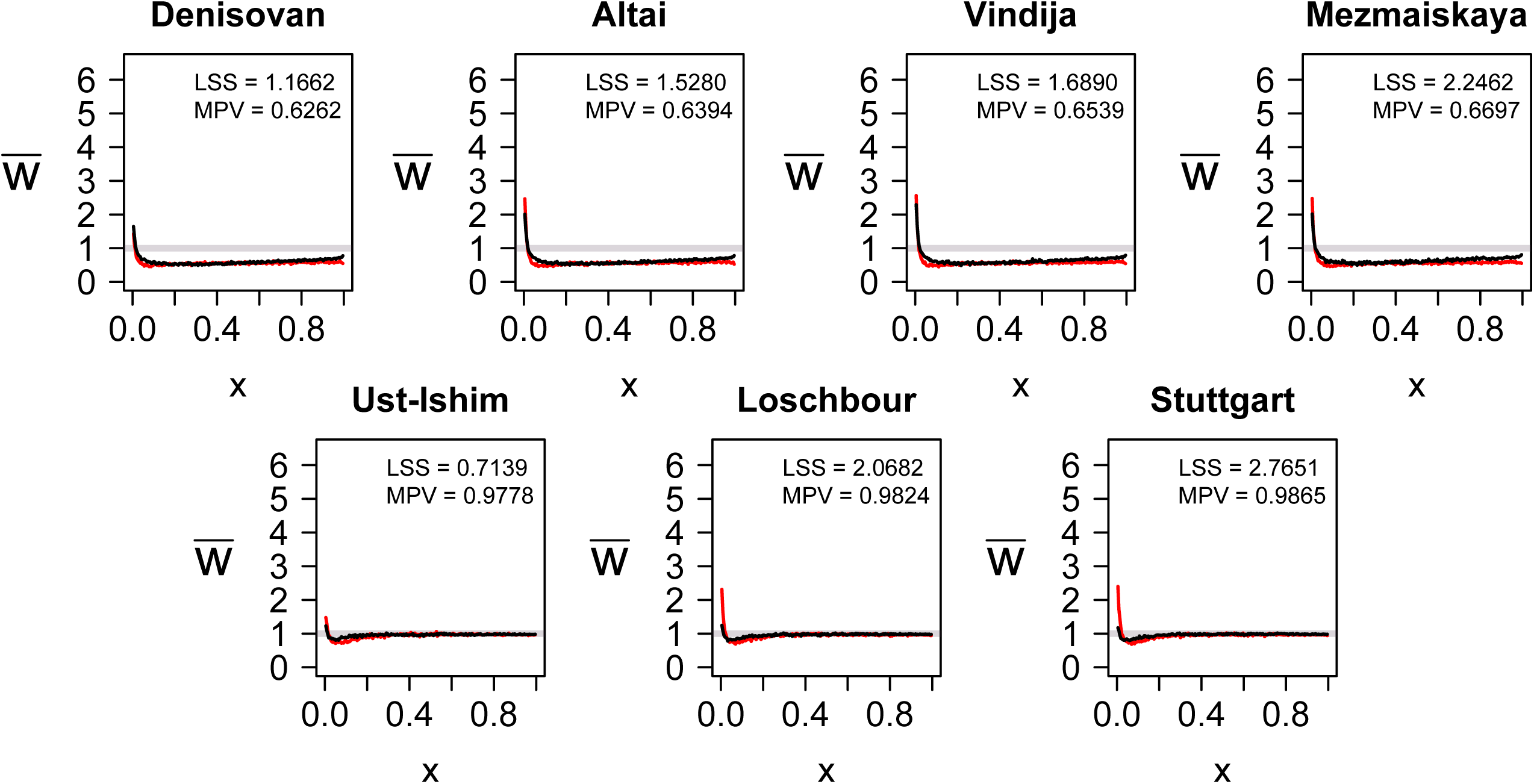
Projection of ancient hominin genomes onto the Yoruba reference panel, refYRI (black), and simulated projection (red). The sum of least squares (LSS) score gives the fit between the observed and simulated projections. The mean projection value (MPV) is the mean for *x* > 0.5.

For the refCHB reference panel (Figure 5), the projections for the Neanderthal and Denisovan genomes are nearly identical to that observed for the refCEU panel (mean MPV = 0.4288, sd = 0.0154). The Ust-Ishim, Loschbour and Stuttgart projections all indicate they are not members of the Han Chinese population (MPV = 0.8626, 0.8748, 0.8632, respectively). Finally, for the refYRI panel, the projections are unusual, but similar to that observed by Yang et al. (2014). The Neanderthals and the Denisovan have a lower projection onto the refYRI panel (mean MPV = 0.6473, sd = 0.0187, Figure 6) than the non-Africans. The higher mean MPV is probably because the Yoruba did not undergo the same bottleneck detected in non-Africans. For non-Africans, the projection increases for common alleles, which was shown in simulations in Yang et al. (2014) to be due to high levels of ancient admixture between the ancestral Yoruba and non-African populations, as well as a population decline in the Yoruba population. Given that the three ancient human samples are all non-African, it is expected that they should all have projections very similar to those observed for modern non-Africans relative to the refYRI panel (Yang et al. 2014).

## COMPARING THE PROJECTIONS TO A SIMULATED DEMOGRAPHY

To gain greater perspective on how the projections of these ancient genomes relate to human demographic history, we compared the ancient genomes to simulated projections taken from a proposed demographic model. We used the demographic model that best fit the set of projections for modern humans published in Yang et al. (2014), which included eight populations of European, African, East Asian and Papuan origin, and the Altai Neanderthal and Denisovan. For each ancient genome, we simulated the same demographic model, adding a single simulated sample retrieved at the time indicated in Table 1, where one generation is assumed to be 25 years. The Neanderthals were placed on the Neanderthal lineage, the Denisovan on the Denisovan lineage, the Ust-Ishim genome shared a common ancestor with Europeans and East Asians, and the Loschbour and Stuttgart genomes were placed on the European lineage, in accordance with the conclusions of their respective studies (Green et al. 2010, Meyer et al. 2012, PrÜfer et al. 2014, Fu et al. 2014 and Lazaridis et al. 2014).

Using *fastsimcoal2* (ver 2.1, Excoffier et al. 2013) and Brent’s algorithm, the time and amount of Neanderthal admixture, the time of Neanderthal divergence and the recent admixture from Europeans to Yoruba were allowed to vary to improve the fit of the projections. The sum of least squares (LSS) was calculated when each simulated and real projection was compared. Using a time of Neanderthal/Denisovan divergence of 610,175 yrs, with admixture into non-Africans 38,950 yrs ago of 0.018, and recent admixture 7,500 years ago from Europeans to Africans of 0.02, the simulated projections were very close to the actual projections (Figure 4-6, LSS score, topright corner).

## DISCUSSION

Simulated scenarios show that the projection can distinguish between samples directly ancestral to a reference population and samples that belong to a sister population that diverged from the reference population. The projections of the Neanderthals and Denisovans all show a very similar projection to each other with respect to each reference panel, despite the differences in sampling time. Therefore, these genomes belong to a sister group and the reconstructed demographic history that recovers the observed projections also places them all in a sister group. These results concur with the conclusions of previous studies (PrÜfer et al. 2014, Meyer et al. 2012, Reich et al. 2010) The increase in rare alleles for their projections onto the refYRI panel was recovered by including some recent admixture from Europeans to the Yoruba population. Another scenario that was not illustrated here is direct admixture from Neanderthals or a sister group to Neanderthals directly into the ancestral Yoruba population. This is unlikely, as two recent studies have proposed recent admixture from non-African to African populations (Wang et al. 2013, Wall et al. 2013). While we simulated direct admixture from Europeans to the Yoruba, the admixture may have come from a different non-African population, which may improve the fit of the Loschbour and Stuttgart projections onto the refYRI panel.

The Ust-Ishim genome is different from both the European and East Asian panels, showing it is likely not a member of either population, but it behaves similarly to other non-Africans with respect to the Yoruba panel. When a simulated ancient sample was placed directly ancestral to Europeans and East Asians 45 kya, the simulated projection was very similar to the observed projection, illustrating that the shape in the projection can largely be attributed to the population size changes in Europeans and East Asians after the Ust-Ishim was sampled.

The Loschbour and Stuttgart genomes sit on the *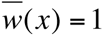* line when projected onto the refCEU panel, but not when projected onto the refCHB or refYRI panel. The projections shows that the Loschbour and Stuttgart could be considered the same population as present day Europeans. In fact, when they were included in the demographic scenario on the European branch, their simulated projections were a very good fit to the observed projections. Determining the projection for these two genomes shows how the projection can distinguish to which populations the test genome belongs or is directly ancestral.

Projections provide a visually appealing method of comparing a single genome against a set of genomes belonging to a well studied reference population. When genomes sampled are ancient, the projection can distinguish between several different demographic scenarios, providing further insight into potential demographic models to test in further, more statistically rigorous analyses. Here, we have shown the projection results for several ancient hominin genomes, but the projection can be applied to any number of organisms.

## CONCLUSIONS

Projection analysis is a useful tool for gaining a rough indication of the demographic history between two populations. Here, we have demonstrated the effects on the projection when ancient samples are included. For scenarios where the ancient population is directly ancestral to the modern population, if the test genome is ancient and the reference panel is modern, the projection reflects the changes in the reference panel since the sampling time. However, when the test genome is modern and the reference panel is ancient, the test genome looks like a member of the reference population.

In the alternate scenario where the ancient population is a member of a sister population, if the test genome is ancient and the reference panel is modern, the projection looks the same as when the test genome is sampled from the present. In the reverse situation when the test genome is modern and the reference panel is ancient, the projection of the test genome moves closer to the 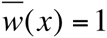 as the reference panel is nearer to the time of divergence.

We studied the projections of several ancient hominin genomes. Neanderthals and Denisovans were not directly ancestral to modern humans. The Ust-Ishim projection looks ancestral to both Europeans and East Asians, and the Loschbour and Stuttgart projections suggest that they are ancestral to Europeans, but not to East Asians or the Yoruba.

Projections provide insight on the ancestry of the ancient genome and their relationship to present day populations. Future studies of ancient genomes may find projections useful as a test for the ancestral relationship between the ancient sample and present day populations. While not a method of demographic inference, the projection’s shape provides early clues as to the direction of further model testing using formal demographic inference tools, such as *dadi* or *fastsimcoal2*.

## ACKNOWLEDGMENTS

We would like to thank F. Racimo, M. Martin, N. Duforet-Frebourg. and R. Rogers for useful discussion. M.A.Y. was supported by a National Science Foundation Graduate Research Fellowship. M.S. was supported in part by the NIH grant R01-GM40282.

## LITERATURE CITED

The 1000 Genomes Project Consortium, G. A. McVean, Altshuler D. M., Durbin, R. M., Abecasis, G. R., D. R. Bentley et al., 2012. An integrated map of genetic variation from 1,092 human genoems. Nature 491: 56–65.

Excoffier, L., Dupanloup I., Huerta-Sanchez E., Sousa V. C. and M. Foll. 2013 Robust demographic inference from genomic and SNP data. PLoS Genet 9: e1003905.

Fu, Q., Li, H., Moorjani, P., Jay, F., S. M. Slepchenko et al. 2014. Genome sequence of a 45,000-year-old modern human from western Siberia. Nature 514: 445–449.

Green, R.E., Krause, J., Briggs, A.W., Maricic, T., U. Stenzel et al. 2010. A draft sequence of the Neandertal genome. Science 328:710–722.

Gutenkunst, R. N., Hernandez R. D., Williamson S. H. and C. D. Bustamante. 2009. Inferring the joint demographic history of multiple populations from multidimensional SNP frequency data. PLoS Genetics 5.

Lazaridis, I., Patterson, N., Mittnik, A., Renaud, G., S. Mallick et al. 2014. Ancient human genomes suggest three ancestral populations for present-day Europeans. Nature 513: 409–413.

Meyer, M., Kircher M., Gansauge M. T., Li H., F. Racimo et al. 2012 A high-coverage genome sequence from an archaic Denisovan individual. Science 338:222–226.

Prüfer, K., Racimo F., Patterson N., Jay F., S. Sankararaman et al. 2014 The complete genome sequence of a Neanderthal from the Altai Mountains. Nature 505:43–49.

Reich, D., Green, R. E., Kircher, M., Krause, J., N. Patterson et al. 2010. Genetic history of an archaic hominin group from Denisova Cave in Siberia. Nature 468:1053– 1060.

Wang, S., Lachance, J., Tishkoff, S. A., Hey, J., and J. Xing. 2013. Apparent variation in Neanderthal admixture among African populations is consistent with gene flow from non-African populations. Genome Biol Evol 5:2075–2081.

Wall, J. D., Yang, M. A., Jay, F., Kim, S. K., E. Y. Durand et al. 2013. Higher levels of Neanderthal ancestry in East Asians than in Europeans. Genetics 194:199–209.

Yang, M.A., Harris, K., and M. Slatkin, 2014. The projection of a test genome onto a reference population and applications to humans and archaic hominins. Genetics 198:1655–1670.

